# Strehler-Mildvan correlation is a degenerate manifold of Gompertz fit

**DOI:** 10.1101/064477

**Authors:** Andrei E. Tarkhov, Leonid I. Menshikov, Peter O. Fedichev

## Abstract

Gompertz empirical law of mortality is often used in practical research to parametrize survival fraction as a function of age with the help of just two quantities: the Initial Mortality Rate (IMR) and the Gompertz exponent, inversely proportional to the Mortality Rate Doubling Time (MRDT). The IMR is often found to be inversely related to the Gompertz exponent, which is the dependence commonly referred to as Strehler-Mildvan (SM) correlation. In this paper, we address fundamental uncertainties of the Gompertz parameters inference from experimental Kaplan-Meier plots and show, that a least squares fit often leads to an ill-defined non-linear optimization problem, which is extremely sensitive to sampling errors and the smallest systematic demographic variations. Therefore, an analysis of consequent repeats of the same experiments in the same biological conditions yields the whole degenerate manifold of possible Gompertz parameters. We find that whenever the average lifespan of species greatly exceeds MRDT, small random variations in the survival records produce large deviations in the identified Gompertz parameters along the line, corresponding to the set of all possible IMR and MRDT values, roughly compatible with the properly determined value of average lifespan in experiment. The best fit parameters in this case turn out to be related by a form of SM correlation. Therefore, we have to conclude that the combined property, such as the average lifespan in the group, rather than IMR and MRDT values separately, may often only be reliably determined via experiments, even in a perfectly homogeneous animal cohort due to its finite size and/or low age-sampling frequency, typical for modern high-throughput settings. We support our findings with careful analysis of experimental survival records obtained in cohorts of *C. elegans* of different sizes, in control groups and under the influence of experimental therapies or environmental conditions. We argue that since, SM correlation may show up as a consequence of the fitting degeneracy, its appearance is not limited to homogeneous cohorts. In fact, the problem persists even beyond the simple Gompertz mortality law. We show that the same degeneracy occurs exactly in the same way, if a more advanced Gompertz-Makeham aging model is employed to improve the modeling. We explain how SM type of relation between the demographic parameters may still be observed even in extremely large cohorts with immense statistical power, such as in human census datasets, provided that systematic historical changes are weak in nature and lead to a gradual change in the mean lifespan.

## 1. Introduction

In most animals, including humans, aging leads to exponential increase of mortality *M*(*t*) = *M*_0_ exp(*αt*) where *t* is age, the dependence commonly referred to as the Gompertz law [4]. It has been long assumed, based on both empirical evidence and theoretical arguments, that the Initial Mortality Rate, *M*_0_ or IMR, and the Gompertz exponent slope, *α*, are universally related by Strehler-Mildvan (SM) correlation

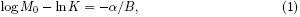

where *K* and *B* are constants introduced in [9], and *α* is inversely proportional to the Mortality Rate Doubling Time (MRDT). SM correlation is not a trivial fact and may not be universal. For instance, it is inconsistent with a recently observed scaling law in [10], which implies that the surviving fractions of *C. elegans* worms under different environmental conditions or mutants with different lifespans can be cast into a universal function of a properly rescaled dimensionless age. This is indeed remarkable, since the experiment establishes proportionality between IMR and *α*, rather than that of log *M*_0_ and *α*, as would be predicted by Strehler and Mildvan. To make sense of the apparent contradiction, we reviewed the problem of Gompertz parameters inference from experimental survival fraction curves (lifetables).

We started by addressing the performance of least squares fit method and immediately found that it may easily lead to an ill-defined non-linear optimization problem, which is extremely sensitive to sampling errors and the smallest systematic demographic variations. We found that whenever the average lifespan of species greatly exceeds MRDT, small variations in survival records are amplified and produce large deviations in the identified Gompertz parameters along the line, corresponding to the set of all possible IMR and MRDT values, roughly compatible with the properly determined value of an average lifespan in the experiment. The best fit estimates in this case turn out to be related by a form of SM correlation. Therefore, we have to conclude that under most common circumstances combined property, such as the average lifespan, rather than IMR and MRDT values separately, can only be reliably determined in experiments with a homogeneous animal cohort due to its finite size and/or a low age-sampling frequency, typical for modern high-throughput settings.

We support our theoretical findings by careful analysis of experimental survival records obtained in a recent series of experiments with *C. elegans* under the influence of lifespan modifying treatments or environmental conditions [13, 10]. The data provides a unique opportunity since, in contrast to similar human mortality records, the laboratory experiments are well controlled for genetic and environmental variations. We start by sampling the worms life histories from the control groups into cohorts of various size and show that the immediately apparent SM correlation in such perfectly homogeneous cohorts is the direct consequence of the hyper-sensitivity of the least squares fit procedure exacerbated by a large sampling error due to insufficiently large number of the animals in the cohorts. In agreement with our theoretical considerations, we find that it is easy to produce a reliable estimate for the average lifespan of such small and homogeneous cohorts, yet it is considerably more difficult to yield reliable estimates of the Gompertz parameters unless a Kaplan-Meier plot is extremely smooth to infer a unique solution.

We conclude observing, that since SM correlation may show up as a consequence of the Gompertz fit degeneracy, which is closely related to the extreme exponential nature of mortality, its appearance may not be limited to homogeneous cohorts only. We show that the same degeneracy along with the extreme dependence of the parameter estimates remains to be an issue with more advanced, such as e.g. the Gompertz-Makeham, models. Finally, we explain how the SM type of relation between the demographic parameters may persist even in extremely large cohorts with immense statistical power, such as in human census datasets, provided that the systematic historical changes are weak in nature and lead to a gradual change in the mean lifespan.

## 2. Strehler-Mildvan correlation is a degenerate manifold of Gompertz fit

To investigate the influence of a factor, such as therapy or a mutation, on aging, one may want to estimate the effects of the experimental design conditions on aging model parameters, such as, for example, commonly used quantities *M*_0_ and *α* of Gompertz law. A natural way to achieve the goal is to analyze Kaplan-Meier plots and fit an experimentally observed fraction of animals surviving by age *t*, *N*(*t*), onto the model prediction. According to Gompertz law, mortality at a given age, *t*, is the exponential function of age, *M*(*t*) = *M*_0_ exp (*αt*). The fraction of the animals alive by the same age is given by the expression 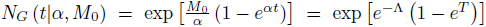, where *T* = *αt* is the dimensionless age and the Gompertz logarithm or curvature, Λ = log (*α*/*M*_0_) = log (1/*M*), is expected to be large. For human patients, for example, we have 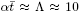 and hence the average lifespan, 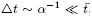, greatly exceeds the MRDT. In this case survival fraction as a function of age drops from 1 to 0 in a short age interval, 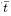, around the mean lifespan 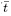, so that 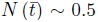 and 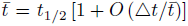. Hereinafter we refer to this parameter range as the extreme Gompertizian limit. A typical behavior of the survival fraction is qualitatively depicted in Fig. 1A.

**Figure. 1:**
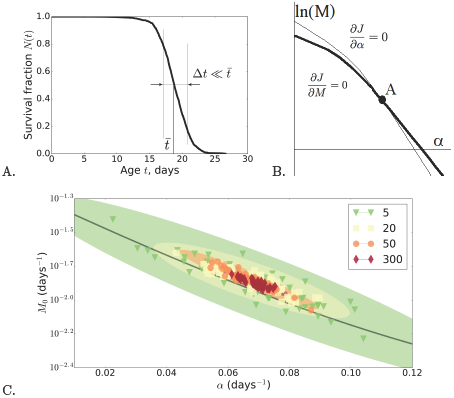
A. Kaplan-Meier survival fraction for a wild-type control cohort from Stroustrup et al. [10]. The function drops from 1 to 0 in a short age-interval, ∆*t*, usually considerably shorter than the average lifespan 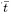; B. A graphic solution of Eqs. (3) in ln *M* and *α* plain. The thick and the thin lines correspond to the vanishing derivatives of the objective function *J* with respect to the parameters *M* and *α*, see (Eq. (3); C. The Gompertz parameters *M*_0_ and *α* obtained from the least-squares fit (2) for the sub-groups of the wild-type control animals (the data from 768 control cohorts from [13]). Here the number of randomly chosen cohorts in sub-groups is *N_coh_* = 5, 20, 50 and 300 (approximately 10 worms in each cohort). The colours mark the numbers of cohorts. For each randomly selected sub-group, a Kaplan-Meier survival curve is obtained and a random realization of the Gompertz parameters is defined from it. Each dot represents a fitting result obtained using data from a particular randomly selected sub-group of the wild-type control animals. Black line represents the iso-average-lifespan curve for the average lifespan calculated for all wild-type control experiments. The variations in determination of *M*_0_ and *α* for different random selections of sub-groups allow us to estimate their fitting uncertainty, which is represented graphically by the characteristic guide-to-eye ellipses in *M*_0_ versus *α* plane.

One way to fit an experimentally observed dependence *N*(*t*) to its Gompertz law estimate is to minimize the mean squared error between expected and observed values of the surviving fraction. Normally, the values of *N*(*t*) are known at a series of discrete age-points. To simplify the analysis we will assume that the number of the observation and the experimental cohorts size are large enough, so that the discrete set of observations *N*(*t*) can be viewed as consequent measurements of a smooth function of age. Under the circumstances, the objective function can be represented as the integral

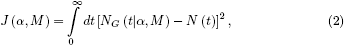

instead of the sum over all observations, to be minimized with respect to the values of *α* and *M*. The procedure can be understood as a standard log-likelihood minimization. The full analysis of the optimization problem is presented in Appendix A. Let us summarize briefly in this section the most important results for the discussion below.

First of all, the best fit Gompertz parameters are solutions of the system of equations

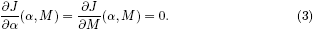

A graphical solution is represented in Fig. 1B. The slopes of the lines, representing the solutions of Eqs. (3) at the intersection point differ by a mere factor *C*_2_/Λ^2^, with *C*_2_ =*O*(1) being a constant. For sufficiently long lived species, Λ ≫ 1, hence the solutions are practically indistinguishable from each other near the intersection point, defining the extremum.

The near collinearity of the directions defined by Eqs. (3) is established in Appendix A, is a consequence of the step-wise nature of the Gompertz survival curve and thus is a signature of a degeneracy of the original inference problem. In Appendix B we show in sufficient details, that the degeneracy of the minimization problem leads to instability of Gompertz fit estimates: should the observed values of *N*(*t*) change a little, *N*(*t*) → *N*(*t*) + *δN*(*t*), either due to sampling errors or to minor systematic biological effects, the estimated quantities will also change, but very strongly,

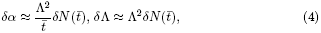

respectively. Remarkably, even though the demographic parameters are subject to a large uncertainty separately, the resulting estimates can only vary in a tightly correlated way

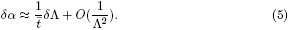

Eqs. (4) and (5) are the central result of the presented study and highlight extreme sensitivity of the Gompertz fit to the slightest variations of the survival fraction in Eq. (2). Therefore, instead of a single and well-defined solution, under most common circumstances one rather gets the whole degenerate manifold of possible values, even in the most idealistic case, when the biological conditions are exactly the same for all experimental replicates and the animal groups under investigation are biologically homogeneous. Any minor survival curve variations lead to the estimates deviations, which are strongly amplified along the line in the (*α*, Λ) plane, defined by Eq. (5) close to the pair of the optimal values, while the fluctuations of the determined parameters were greatly suppressed in the orthogonal direction. The appearance of the SM correlation and the instability of Gompertz parameters estimates are demonstrated by numerical analysis of simulated cohorts of different size, see Fig. A.4 in the Appendix.

The established degeneracy of the fit has a surprising and unintended consequence. Let us assume that the experiments are repeated in exactly the same conditions. Then the sampling error is the only source of the uncertainty in the survival records, and, therefore, each of the possible fits corresponds to the correct value of the average lifespan of the demographically homogeneous cohort, *t_exp_* = ∫ *dtN*(*t*). Since, in this case, 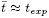 is the same in every calculation, Eq. (5) can be integrated to yield

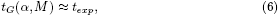

where *t_G_*(*α*, *M*) is the Gompertz law prediction for the mean lifespan. This means that every pair of *M* = *M*_0_/*α* and *α* values, satisfying Eq. (6) can, in principle, be obtained from the analysis of different realizations of the same experiment. To support our conclusion, we checked this sensitivity in a numerical simulation (Fig. A.4). Since *t_exp_* ≈ Λ/*α* in the Gompertz model in Λ ≫ 1 limit, the degeneracy manifold is approximately defined by log *M*_0_ — log *α* = —*αt_exp_*, which is nothing else but a form of SM correlation, defined by Eq. (1)! We propose that the “canonical” form of SM relation (1) can be viewed as an approximation to the “exact” form (6) in the extreme Gompertz limit, Λ ≫ 1. The restriction on Λ is totally unnecessary, although it leads to a more tractable form of Eq. (6). A more accurate, but somewhat cumbersome, approximation for the average lifespan *t_G_*(*α*, *M*) from the Gompertz law for an arbitrary value of Λ and therefore a more precise form of the expected SM correlation in homogeneous cohorts is derived in the appendix, see Eq. (C.1).

## 3. Gompertz fit examples in homogeneous populations

To support our theoretical arguments and back up the conclusions below, we turn to a set of relevant experimental data. We choose to investigate the Kaplan-Meier plots from two recent high-throughput experiments, addressing effects of a series of pro- [13] and anti- [10] longevity environments or interventions. All the data are obtained in the fully controllable settings in the studies with genetically homogeneous cohorts. The level of control is, of course, totally unmatched in any kind of studies of human mortality.

The first data set [13] describes life histories of 1416 experimental cohorts of the same animals, living in various laboratory conditions. Typically, each of the cohorts consists of about 10 worms. A few of the experiments were repeated to achieve higher significance. 768 or almost half of the cohorts are control ones. We used the data to estimate uncertainty of Gompertz fit parameters in the ideal case of a genetically homogeneous population in identical experimental conditions in a particular laboratory. The data let us simulate results of Gompertz fit in a typical demographic study of an aging population. To do this, we choose a subset of *N_coh_* cohorts at random from all 768 available aging control cohorts. We used *N_coh_* = 5,20,50 and 300 and calculated the Kaplan-Meier survival plots each time using a subset of the animals and fed it to the least squares fit. The procedure was repeated enough times to get a reliable estimate of the fitting parameters uncertainty for every chosen value of *N_coh_*. The results of the computational experiment are presented in Fig. 1C. Every simulated cohort is represented by a dot in the (*α*, log *M*_0_) plain. The uncertainty of the estimates is larger for smaller cohorts. To support our conclusion, the obtained result was checked on a numerically simulated data set (Fig. A.4, Appendix A). This is totally expected, if the sampling error is due to the finite number of animals in the group and is therefore smaller in the larger groups. Even though Gompertz logarithm Λ ≈ 1.3 is not particularly large, the uncertainty of the fit is amplified along SM correlation line, according to Eq. (5). Therefore, we are able to show that SM correlation may show up even under such an ideal study protocol, when by definition there is no underlying biological difference between the subsequent repeats of the same experiment.

In addition to the life histories of control cohorts, the data set provides information on life-prolonging effects of a large number of currently used and experimental drugs. We selected the cohorts under the treatments, with the largest number of experimental replicates and, hence, with the least expected degeneracy of the Gompertz parameters. We can assume that for each of the treatments the corresponding cohorts are homogeneous, since all other experimental and genetic conditions are controllable and are exactly the same in all the replicates. We estimated the effects of the fitting degeneracy by selecting 2/3 of the animals in each treatment group at random, and producing Gompertz parameters from the corresponding simulated Kaplan-Mayer plots. The procedure lets us estimate the uncertainty of the fit and represent it graphically by the characteristic ellipses in *M*_0_ versus *α* plane in Fig. 2. To compare, we also plotted a red black-edged ellipse corresponding to the same kind of estimate for the largest control group, corresponding to *N_coh_* = 300 (~ 3000 animals). Remarkably, the degeneracy of the fit is still pretty profound, even for such a reasonably large group. We estimate the relative error in the determination of Gompertz parameters in the control groups to be about 15%, and hence the Gompertz exponent estimate *α_wt_* = 0.07 ± 0.01 days^−1^ for the wild type animals. The iso-average-lifespan curves (SM correlation curves) are plotted and coloured according to the average lifespan in the cohorts as shown in the inset. In agreement with Eq. (4), the calculations demonstrate, using the real world data, that small fluctuations of a survival curve even for homogeneous cohorts, due to a finite number of animals in them or a low age-sampling rate, lead to the strong fluctuations of the fitting parameters along SM curves, whereas fluctuations orthogonal to these SM curves are substantially suppressed.

**Figure. 2:**
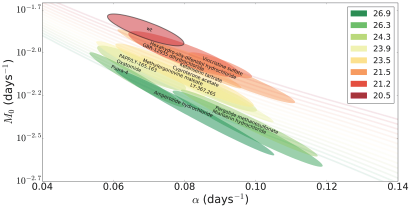
IMR (*M*_0_) versus Gompertz exponent slope *α* estimates obtained with the least-squares fitting procedure using lifetables from the prolongevity *C. elegans* experiment [13]. To obtain uncertainty of the Gompertz parameters (depicted by the sizes of ellipses) for each of these drug-treatments, the same random selection procedure as in Fig. 1C was employed, but for each drug-treatment 2/3 of all the available cohorts (roughly 20 cohorts, 10 worms in each) were randomly selected to calculate a particular random realization of Kaplan-Meier plot and the corresponding random realizations for the Gompertz parameters. The coloured solid lines correspond to analytical, asymptotic expressions representing the iso-average-lifespan manifolds defined by Eq. (C.2), where the colours mark the corresponding average lifespans. The position and size of the red black-edged ellipse indicate the best estimates for the Gompertz fit parameters and their uncertainty for the control wild-type experiment with 300 cohorts (~ 10 worms in each), which corresponds to the red dots in Fig. 1C. The coloured ellipses (from green to red) represent typical Gompertz parameters with their uncertainty for pro-longevity treatments [13], each of them corresponds to a particular drug-treatment experiment with roughly 30 cohorts (~ 10 worms in each) on average which is 10 times smaller than for the wild-type control case with 300 cohorts of roughly 10 worms in each, thus it leads to a much larger uncertainty in Gompertz parameters estimation (larger ellipses).

Life histories from the experiment reported in [10], represent another interesting limit, since death events are now sampled at a very high frequency (almost every second for several thousand animals in a single cohort), and therefore the resulting Kaplan-Meier survival plots are extremely smooth and Gompertz parameter estimates are perfectly reliable. Figs. 3A and B summarize analysis of the two experiments, [13] and [10]; all the data points are plotted on the same graphs and recoloured according to the new global age-pseudocolour scale. Once again, the figures demonstrate that Gompertz parameters determination uncertainty in every homogeneous cohort manifests itself as fluctuations amplified along the iso-average-lifespan curves, although the error margins are now much smaller due to a dramatically higher precision of the Kaplan-Meier plots. IMR and Gompertz slope values estimated from [10] appear to be very reliable representations of the underlying biology. Therefore, independence of IMR from *α*, is significant and hence is the experimental proof of a very non-SM-like relation between the parameters.

## 4. Conclusions and Discussion

Throughout the paper we employ the least squares fit of survival fraction in the form of Eq. (2) as a practical and mathematically tractable way of the Gompertz parameters inference. A commonly accepted alternative is to produce the fit using the log-transformed values of empirical mortality rate in a data set. The latter possibility looks more attractive from a conceptual viewpoint, but its practical applications may be hindered by the following considerations. The mortality rate needs to be estimated by numerical differentiation of experimental survival lifetables, which are by no means continuous and differentiable, especially late in life. On top of that, the empirical mortality does not necessarily follow Gompertz law. For example, the risk of death may grow exponentially first and then saturate at a constant level at extreme ages (phenomenon known as mortality deceleration, see e.g. [11]). Whenever this happens to be the case in the data, an extra assumption is required to specify the age range, where Gompertz fit is to be performed. Normally, one restricts the analysis to a reasonable vicinity of age, corresponding to average lifespan of animals in the experiment. The procedure suggested here may not be necessarily better, but it is easier to understand using analytical tools, requires only raw experimental data points, a survival curve, and produces a unique answer without extra assumptions. In Appendix A we show that minimization of Eq. (2) yields the best Gompertz approximation of the data, using the data points mostly close to the average lifespan in the cohort, and therefore is consistent with the “standard” definition, whenever the Gompertz fit works well.

The immediate and somewhat surprising conclusion from the presented study is that even the two-parametric Gompertz fit may easily turn out to be an ill-defined mathematical procedure. Not to mention proportional hazards models and accelerated failure time models with quite a few extra variables, which might obfuscate researchers with miscellaneous degenerate manifolds of their numerous parameters. We demonstrated that a form of SM correlation occurs every time when the survival fraction is changed, even by a small bit, either due to sampling errors or true biological reasons. The coordinated change of IMR and *α* values, amplified along the line of SM correlation, corresponding to the expected value of the mean experimental lifespan, is a property of a fit involving Gompertz survival function in the extreme Gompertz limit, when the mean lifespan exceeds MRDT. This should be, of course, especially true in human demographic data analysis, where SM correlation was identified first in [9]. The lifespans in the cohorts representing the countries in the original work were pretty close, 59 ± 9 and 62 ± 9 years for males and females, respectively, not to mention that the set of countries was rather unrepresentative, since the period mortality data were restricted to a very narrow time interval 1947 – 53 for 32 relatively well-developed countries. Nevertheless, Gompertz parameter estimations ended up distributed in a vast range of *α* = 0.08 ± 0.04 years^−1^ and exhibiting a very high degree of correlation between *α* and log *M*_0_.We believe that the dependence is then nothing but the narrow stripe enclosing a degenerate manifold of Gompertz parameters corresponding to the average lifespans.

**Figure. 3:**
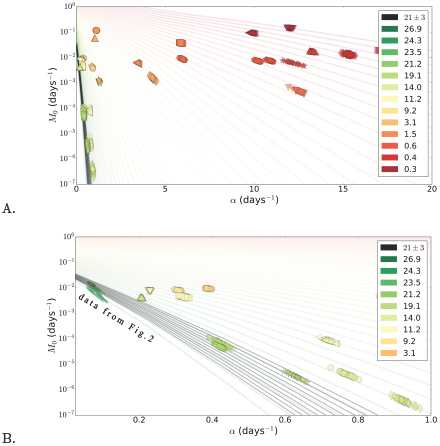
A. A combined IMR (*M*_0_) versus the Gompertz exponent slope *α* plot, summarizing the estimates obtained by the least-squares procedure using *C. elegans* lifetables from experiments [13, 10]. The colours denote the average lifespans in days. Each pair (colour, marker) corresponds to a single realisation of an experimental cohort, to estimate variation of Gompertz parameters inside the homogeneous cohorts, the replicates are obtained by the same reshuffling procedure as for Fig. 1 and 2. The coloured curves are analytical asymptotic expressions corresponding to the iso-average-lifespan manifolds defined by Eq. (C.2). The black curves are iso-average-lifespan curves within a narrow interval of lifespans 21 ± 3 days B. A zoom into *α* ε [0,1] interval in Fig. 3A. The points from Fig. 2 are also shown, recoloured according to the new age-pseudocolour scale.

A possibility of superficial nature of SM correlation has been already highlighted by L. and N. Gavrilovs in [2, 3], where a more sophisticated aging model, Gompertz-Makeham version of the mortality law, was invoked to produce a more stable fit. In Appendix D we show that inclusion of Makeham term into the model may help fit the experimental data in a different aging regime with a significant age-independent component in mortality rate, whenever it is required. However, we observe that the more advanced version of the survival model cannot help improve considerably the fit performance in a situation, when an experimental Kaplan-Meier plot is insufficiently sampled due to a limited number of animals in a cohort or a low age-sampling rate.

Aging in humans is an extreme case, corresponding to Λ ≈ 10 in our calculations. It should thus be not surprising that more than half a century after the publication of the original work [9], the idea of SM correlation and biological picture behind it are deeply rooted in aging studies (see ex. [14, 12, 1, 6, 7] as examples of recent works) but there is still no consensus explanation of its origin. Most notably, heterogeneity of aging populations was recently investigated and proposed to explain SM correlation [12, 14, 5]. For example, [14] observed SM correlation in human mortality data from 1955 to 2003 showed that its parameters evolve over time. This goes well along with our expectations, since we believe that human demography changes rather slowly over time and for every period of several years any human population can be considered nearly homogeneous with a constant lifespan. Therefore, there must exist an approximate form of SM correlation as was shown in the present study. Needless to say, the lifespan and the quality of environmental conditions, such as the available achievements of medicine, progress steadily in nearly all countries and their effect on the shape of survival curve and hence on Gompertz parameters can be observed on a larger timescale as a gradual change of SM correlation parameters. For example, [6] observes a breakdown of the long-term SM trend in the mid-1950s, as a result in the recent years the literature on SM correlation has shifted to the reasoning behind this breakdown. To provide our own interpretation of this phenomenon, we show analytically in Appendix B and via illustrative human mortality analysis in Appendix E how a gradual demographic change produces distortion to the SM correlation plot, which is usually a collection of familiar iso-average-lifespan manifolds, corresponding to practically homogeneous sub-populations born in the interval of several adjacent years with neither appreciable lifespan, nor the outer environment differences. The intra-group differences are greatly amplified along the tangent lines, described by Eq. (5), followed eventually by a fundamental trend in improving quality of life (see Fig. E.5 for the illustration of period female human mortality data analysis for France and Sweden in 1860-2000). We conclude our analysis having observed that the peculiar breakdown of global linear trend in 1950s is probably associated with Makeham age-independent mortality, since there is a 10-fold sharp drop in its value in the vicinity of mid-1950s, which is marked by the sudden colour-change of the ellipses in Fig. E.5 for two parts of the “broken” SM correlation. Human mortality is by no means a two- or three-parameter process, hence all our considerations are limited to Gompertz and Gompertz-Makeham aging regimes, whereas there might exist other higher dimensional changes in mortality for particular time periods, analysis of which is out of scope of the present paper.

A number of theories suggesting some physical or biological background behind SM correlation have been proposed since its discovery. In our work, we come to the conclusion that any biological interpretations should be used cautiously, especially those based on experimental data sets with poor quality of experimental Kaplan-Meier survival plot, due to a limited number of animals in a cohort or a low age-sampling rate. To establish variation in lifespan with sufficient accuracy is not a small feat, requiring large cohorts of aging and dying specimens in an experiment. The analysis leading to Eq. (4) demonstrates that one would need at least a Λ^4^ times larger cohort to produce separate estimates of Gompertz parameters with the same precision. Otherwise, one might misinterpret the iso-average-lifespan manifold in Gompertz parameters plane as a totally spurious SM correlation between Gompertz parameters even in homogeneous cohorts. The presented analysis is very technical in nature, we were only concerned with statistical properties and robustness of Gompertz parameters inference from experimental data. It still may be the case that IMR and MRDT are indeed mechanistically related by nature of underlying biology of aging. We believe that the question can eventually be settled as more high quality lifetables are collected in very well controlled studies and are made available for researchers. We are leaving these considerations for future research.

## Acknowledgements

We thank Prof. Petrascheck, Prof. Fontana and Nicholas Stroustrup for providing raw survival data, and experimental lifetables from [13, 10]. We thank Ivan Molodtsov, Natalia and Leonid Gavrilovs, Aubrey de Grey for helpful discussions, Boris Zhurov for help with visualization software, Evgeny Getmantsev for proofreading. This work was supported by Gero LLC.

### Appendix A SM correlation is a degenerate manifold of Gompertz fit in homogenous cohorts

Let us start with the solution for regression problem in Eq. (2), which can be found from the minimum conditions, *∂J*/*∂α* = 0 and *∂J*/*∂M* = 0, where *M* = *M*_0_/*α*, Λ = log(*α*/*M*_0_) = log(1/*M*). Using the relations *N_G_* = exp (*M* (1 – *e^αt^*)), *∂N_G_*/*∂α* = (*t*/*α*)*∂N_G_*/*∂t*, and *∂N_G_*/*∂M* ≈ (*αM*)^−1^*∂N_G_*/*∂t*, we obtain

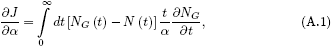

and

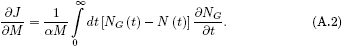

In the extreme Gompertz limit *N* (*t*) and *N_G_* (*t*) are step-like functions and hence we can use the approximations 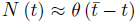 and *N_G_* (*t*) ≈ θ (*t_G_* — *t*). It follows immediately from Eqs. (A.1) and (A.2) that up to terms with a relative error 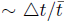, both conditions are equivalent and yield 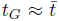. This is a form of correlation between Gompertz fit parameters:

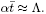

Of course, the equation holds only if Λ = ln(*α*/*M*_0_) ≫ 1. In Appendix C we propose a simple and yet an accurate generalization

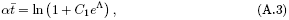

applicable for Λ ~ 1 with *C*_1_ ~ 1 being a numerical constant.

Both expressions in the square brackets, Eqs. (A.1), (A.2) and also the derivative *∂N_G_*/*∂t* behave as sharp peaks located at 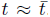. Let us note that next to the intersection point A both curves corresponding to the zero derivatives Eqs. (A.1), (A.2), ί = 1, 2 behave as *α* — *α_A_* ≈ *k_ί_* (*M* — *M_A_*). To estimate the difference between the slopes, it is sufficient to approximate *N*(*t*) by a steep survival function 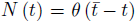. The first equation for the derivative of the objective function with respect to *α* Eq. (A.1) can be rewritten as

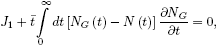

where

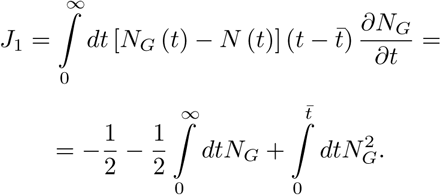

Having collected all terms at most of the order of 1/Λ^2^, we find

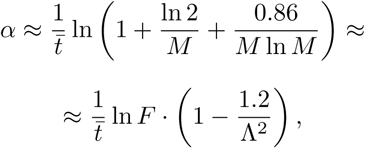

where the approximate relation 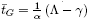 from Appendix C was employed.

The second equation for the derivative of the objective function with respect to *M* Eq. (A.2) takes the form 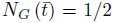 that gives the relation similar to Eq. (A.3):

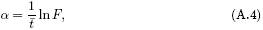

with *F* = 1 + ln(2)/*M*.

Therefore, the two solutions of the optimization problem in Eqs. (A.1), (A.2) are nearly equal and the difference between the slopes at the intersection point is as small as *O*(1/Λ^2^). It means that graphical solutions log*M* (*α*) of Eqs. (3) go very close to each other near an intersection point as qualitatively depicted in Fig. 1B. In Appendix B we show how this degeneracy of the solutions amplifies any small perturbations to the survival function into large fluctuations of Gompertz parameters estimates.

### Appendix B Extreme sensitivity of Gompertz fit estimates with respect to survival plots variations

To see how the Gompertz fit degeneracy in the extreme Gompertz limit of large Λ translates into large uncertainties of inferred Gompertz parameters, let us consider how the model estimates depend on small variations of survival plots. Eq. (A.2) for the derivative of the objective function with respect to *M* can be transformed into

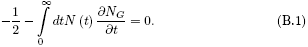

For infinitesimally small changes in the survival fraction Δ*N* (*t*),

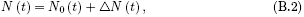

the average lifespan also changes slightly. As usual in this study, we approximate the survival fraction by the step function 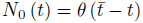 and, from Eq. (B.1), we obtain

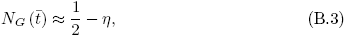

where 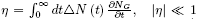. Note that since the function 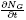 is a narrow peak of the width Δ*t* ~ α^−1^ residing next to the average lifespan 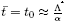, for any slowly varying Δ*N* (*t*) we expect η ≈ Δ*N*(*t*_0_).

**Figure. A.4:**
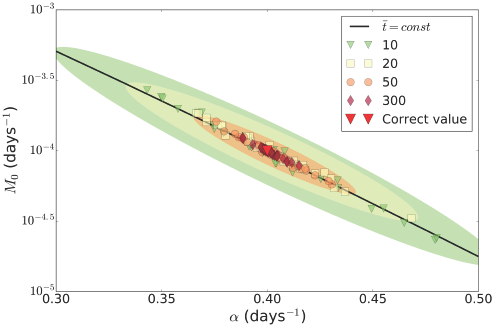
Gompertz parameters *M*_0_ and *α* estimates obtained from the least-squares fit (2) for simulated cohorts with sizes *N* = 10, 20, 50 and 300. The colours differentiate the sizes of cohorts. For each simulated cohort, a Kaplan-Meier survival curve is obtained and a random realization of Gompertz parameters is defined from it. Each dot represents a particular realization. The black line represents the iso-average-lifespan curve for the average lifespan used in this numerical simulation. The variations in determination of *M*_0_ and *α* for different random realizations of noise is represented graphically by the characteristic guide-to-eye ellipses in *M*_0_ versus *α* plane.

From Eq. (B.3) with the help of the approximation (A.3) for the average lifespan we transform the relation *∂J*/*∂M* = 0 into

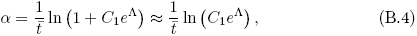

where 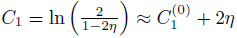 and 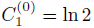. Similarly, the derivative of the objective function with respect to *α* from Eq. (A.1) can be approximately solved as

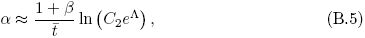

where 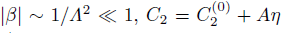 and 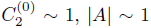 are numerical constants.

To find new values of Gompertz parameters *α* and Λ after the perturbation Δ*N* (*t*) to the survival curve, one should solve Eqs. (B.4), (B.5). The solution can be represented as

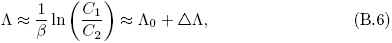

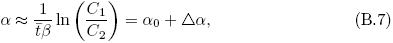

where 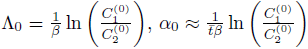 and 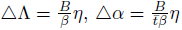 are the “reference” and the perturbed values of the optimization variables. Here |*B*| ~ 1 is another constant factor. Eqs. (4) can be obtained from Eqs. (B.6) and (B.7) using the relation η ≈ ∆*N*(*t*_0_).

From the definition of average lifespan, 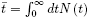, it follows that the average lifespan variation after the perturbation is given by

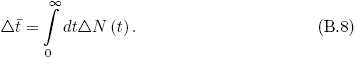

Two important situations should be considered here. Let us study first a consequent realization of the experiment involving homogeneous cohorts. Here *N* (*t*) = *N*_0_(*t*) + Δ*N* (*t*), so that it is natural to assume ∆*N* (*t*) being a random measurement noise, or sampling error, different from one cohort to another. Since 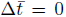 in this case, Eqs. (B.6) and (B.7) immediately yield the linear relation between variations

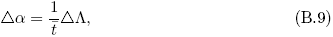

which is the same as Eq. (5) of the manuscript. Importantly, here the sampling error is the only reason for the variations in *α* and Λ. Nevertheless, the inferred values appear to be correlated, the apparent relation coincides with SM correlation. According to Eqs. (4), the errors of Gompertz parameters determination are large, compared to variations in the measurements of average lifespans.

The appearance of SM correlation and the instability of Gompertz parameters estimates can be demonstrated by numerical optimization of the proposed objective function using lifetables of simulated cohorts of different sizes. The results of such a calculation are represented in Fig. A.4. As expected, the sampling error decreases as the cohort size increases. In agreement with Eq. (B.9) the identified Gompertz parameters are distributed in a highly anisotropic way, the direction of maximum variation coincides with SM correlation.

Survival curves may also change systematically in such a manner that the average lifespan is not constant. This can be observed, for example, if the average lifespan is studied in human cohorts collected over sufficiently large periods of time. The human lifespan has been growing slowly over decades and centuries following a fundamental trend in improving quality of life and available medicine. Therefore, one cannot consider such a population as biologically homogeneous on large timescales. Technically, such a systematic change in survival functions produces a net change 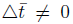, and hence the combined system of Eqs. (B.6), (B.7) and (B.8) should be used to obtain the proper solution. As Gompertz parameters follow variations in biological conditions, the slope 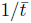 in Eq. (B.9) also changes and hence the point, representing the solution, may depart from the linear SM relation in α and Λ plane.

If lifespan variations are sufficiently small, 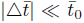, then from Eq. (B.9) it follows immediately that 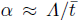 and hence Strehler-Mildvan correlation approximately holds. Conversely, significant changes in lifespan 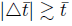 break Strehler-Mildvan correlation, which is clearly the case, for example, in Fig. 3A of the present work, summarizing the Gompertz fit results of the experimental data for *C. elegans* from Stroustrup et al. [10], where the change in the average lifespans is significant enough. It is important to ascertain that the value of 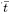 and its modifications are specific to particular experimental conditions. It is impossible to calculate these values explicitly without a rigorous mathematical description of these conditions. In Appendix E we show how a SM-like correlation may show up in a research of human demographics over time in response to gradual decrease of a biological factor, such as age-independent mortality.

### Appendix C Gompertz iso-average-lifespan curves

We have proven above that the only parameter which can be reliably inferred by a gradient descent method in a finite homogeneous cohort is average lifespan.

If mortality in an experimental cohort at hand can be accurately described by Gompertz law, the average life expectancy is

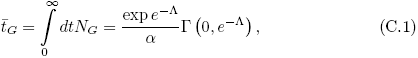

where Λ = ln (*α*/*M*_0_) and г(0, *e*^−Λ^) = *E*_1_(*e*^−Λ^) is the upper incomplete gamma function (or the special function commonly referred to as the exponential integral) which has an expansion

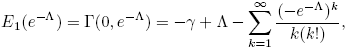

where *γ* ≈ 0.577 is the Euler-Mascheroni constant.

The fit requires that the experimental average lifespan should be equal to the Gompertz average lifespan:

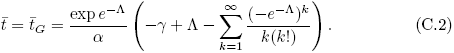

In the extreme Gompertz limit, Λ ≫ 1, the solution is rather simple

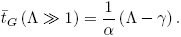

Although the convergence of the series on the right side of Eq. (C.2) is known to be poor for Λ ≤ 1, we still can obtain a divergent series approximation

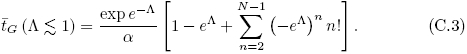

For practical applications, there is a simple and yet an accurate approximation for 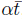, suitable for all possible values of Λ:

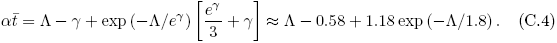

Depending on the value of Λ we obtain: 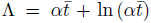 for Λ ≤ 5 and 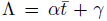 for Λ ≥ 5, respectively. The two asymptotes produce Strehler-Mildvan correlation in the form

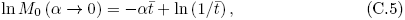

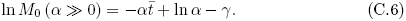

We choose these approximations to plot the correlation curves from Eq. (6) in Fig. 2. More specifically, we put the asymptotic expansions together with a spline at the intersection point 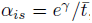, to produce a series of smooth lines. Remarkably, both the exact solutions and the approximations are practically indistinguishable from straight lines.

### Appendix D Gompertz-Makeham iso-average-lifespan curves

Gavrilov [2, 3] proposed that Strehler-Mildvan correlation is a spurious relation in the original work of Strehler and Mildvan [9] due to the neglect of age-independent background mortality. Let us investigate Gompertz-Makeham survival fraction

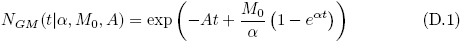

and Gompertz-Makeham fit properties in the same manner as we did for our Gompertz model fit above. The exact average lifespan for Gompertz-Makeham law is

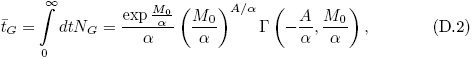

which is indeed somewhat better than its Gompertz law counterpart, since the Makeham parameter moves the centre of our expansion for the upper incomplete gamma function from the singular point near zero argument of 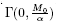 to a better analytical point nearby a non-zero argument of 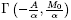. Accordingly, one can expect the divergent series approximation as in Eq. (C.3) to disappear. Nevertheless, we are left with the three-parameter fitting procedure for which SM correlation exists in exactly the same manner. We saw how the degeneracy manifold in case of Gompertz fit arises from the vanishing slope between two optimization curves. In a similar way, in case of Gompertz-Makeham fit the same problem emerges due to the shrinking slope between two optimization surfaces, representing the solutions of Eqs. (3), now in the three-dimensional parameters space. Makeham parameter determines the specific cross-section of these surfaces, where the same degeneracy problem for the remaining pair of Gompertz parameters still persists. Although, we expect that the Makeham parameter A estimate from the fit is well-defined as long as it is large enough. Nevertheless it is not often the case, at least in human aging studies, since the age-independent mortality is usually small and hence the introduction of the Makeham parameter is of a little practical consequence.

### Appendix E An example of Gompertz-Makeham fit for human mortality data

Human mortality data analysis is not of primary interest for the present study, since it is fundamentally difficult to attain genetic and environmental variations control, characteristic to laboratory experiments. For that reason, we provide the following example illustration of how our results concerning the stability of Gompertz fit could be employed for human demography studies. For the demonstration, we have chosen female human period mortality data for France and Sweden [8] collected for the time interval from 1860 to 2000. Since we are now considering lifetables, including those from the time periods prior to 1950, when age-independent mortality was very high, we have to take it into account explicitly and employ Gompertz-Makeham fit.

**Figure. E.5:**
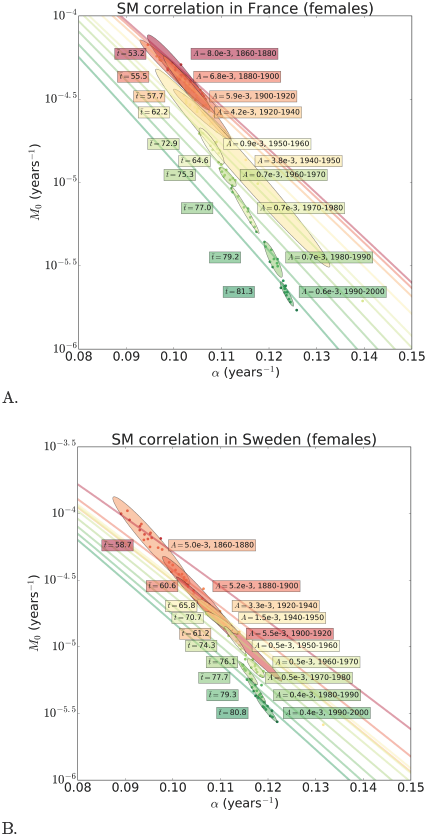
A. A combined IMR (*M*_0_) versus Gompertz exponent slope *α* plot for the female human period mortality data in France from Shkolnikov et al. [8], summarizing the estimates obtained by the least-squares procedure for Gompertz-Makeham mortality law. The dots represent the fitting results for the yearly period mortality data. The dots, the iso-average-lifespan curves defined by Eq. (D.2) and the left boxes with 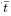 values are coloured according to average lifespan 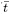. The ellipses combine fitting results for several successive years and show their long-term uncertainty in (*M*_0_, *α*) plane, while the corresponding time periods, average lifespans and Makeham parameters *A* are written in the boxes on both sides of ellipses. The pseudocolour of the right boxes and ellipses denotes the Makeham age-independent mortality value *A*: green for a small and red for a large Makeham contribution. B. The same plot for the Swedish female human period mortality data from Shkolnikov et al. [8].

The results of the calculations for the two countries are summarized in Figures E.5A and B. The fitting results are represented by dots. The solid lines in the background represent iso-average-lifespan curves as prescribed by Eq. (D.2) from Appendix D. Average lifespan is now a function of 3 variables 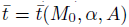. We colour the lines and individual fitting results according to average lifespan. We assumed that underlying biological factors change slowly with time and combined the dots, representing a few subsequent periods into ellipses. Sizes and orientations of the ellipses provide the estimates of fit uncertainties. We coloured the ellipses according to the calculated values of Makeham term in a continuous way, from red to green for large and small age-independent mortality estimates, respectively.

Human demography evolves rather slowly and therefore for sufficiently short time-periods any human population can be considered homogeneous with a constant lifespan. Therefore, we expect that there should exist an approximate form of SM correlation. We indeed observe in Figures E.5A and B all the ellipses aligned more or less along the iso-average-lifespan curves. Not unexpectedly, over longer timescales SM correlation still holds “locally”, but, in line with expectations from Eq. (B.9), the slope of the line changes gradually along with the average lifespan, which increases over the decades, apparently as a function of environmental conditions improvements and achievements of medicine. The breakdown of global linear trend in 1950s is probably associated with a 10-fold Makeham age-independent mortality drop within the first half of the century, which is marked by the sudden colour-change of the ellipses in Figure E.5 between two parts of the “broken” trajectory.

The behavior can be easily understood with the help of the analytical relations derived in Appendix B. Let us assume that Makeham age-independent mortality term is changed by a small margin, *A* → *A* + *Δα*. Then, due to Eq. (D.1) the surviving fraction also changes *N*(*t*) → *N*(*t*)-exp(*—δAt*). Following the derivation leading to Eqs. (B.6), (B.7) and (B.8), we observe that 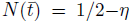, where 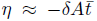 and is presumed to be small. Therefore, while the average lifespan acquires a small correction 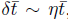, the Gompertz parameters change much stronger, by

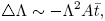

and

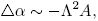

still in the direction of SM correlation, 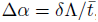, at the given value of average lifespan. The estimates obtained here are interesting on their own. We have just demonstrated, that the Gompertz fit degeneracy does not manifest itself only in obscuring true aging model parameters in a situation when there is an insufficient number of specimens in experimental cohorts. In fact, the spurious correlation between Gompertz parameters may mask a truly biologically significant response to experimental conditions. For example, a therapy aimed at reduction of age-independent mortality, such as an antibiotic, may be mechanistically characterized as an intervention, fundamentally acting on aging by increasing Gompertz exponent and reducing IMR.

